# PepCABO: Latent-space Bayesian Optimization for Peptide-MHC Binding Using Contrastive Alignment

**DOI:** 10.64898/2026.03.13.711540

**Authors:** Mohsen Ghane, Dani Korpela, Alexandru Dumitrescu, Harri Lähdesmäki

**Affiliations:** Department of Computer Science, Aalto University, Espoo, Finland

## Abstract

**Motivation:** Optimizing peptide sequences for binding to specific MHC class I alleles is a central challenge in immunotherapy and vaccine design. The combinatorial size of peptide space, the nonlinear nature of peptide– MHC interactions, and limited experimental budgets make efficient optimization difficult. Latent-space Bayesian optimization (LSBO) provides a framework by embedding discrete sequences into a continuous space where Bayesian optimization can be applied. However, existing LSBO methods do not effectively leverage binding data from related alleles and often rely on inefficient random initialization.

**Results:** We propose PepCABO, an LSBO framework for peptide–MHC binding using contrastive alignment, which utilizes a dual variational autoencoder framework that jointly learns peptide–allele alignment and a Gaussian process surrogate prior to Bayesian optimization. This simultaneous training induces a latent geometry that reflects the binding landscape and enables structured knowledge transfer across alleles. The pretrained model shapes a structured latent space in which peptides with high objective values regarding a specific MHC allele are geometrically organized, while the jointly trained Gaussian process defines an informative prior over the objective in this space, enabling principled and efficient exploration of promising regions during subsequent optimization. Across 12 target alleles without prior binding data and under both low- and high-budget settings, PepCABO consistently outperforms various baselines. We observe faster convergence, improved area under the optimization curve, and stronger best-found binding affinities, suggesting improved sample efficiency under experimentally constrained scenarios.

**Code availability:** The source code is available at https://github.com/mohsen-g/PepCABO

## 1. Introduction

Binding of antigenic peptides to MHC class I molecules is an essential step in cellular immune surveillance, enabling nucleated cells to display intracellular peptides for detection by CD8^+^ cytotoxic T lymphocytes [Wieczorek et al., 2017]. Consequently, optimizing peptide sequences for binding specific MHC alleles represents a major opportunity in immunotherapy and vaccine development [Comber and Philip, 2014]. Despite its importance, this optimization remains a difficult experimental and computational task. First, the peptide optimization space grows exponentially by sequence length, resulting in an astronomical peptide space, which cannot be searched exhaustively. Second, the relationship between peptide sequence and binding affinity is highly non-linear and context-dependent, making accurate prediction of quantitative binding affinity for a given peptide-allele pair a difficult task [Collesano et al., 2024], while experimental measurement of this quantity remains expensive and low-throughput [Haj et al., 2020]. This limits both the reliability of optimization frameworks that rely solely on predictive models and the number of optimization iterations that can be performed using real-world experimental evaluations.

Recent advances in generative modeling and Bayesian optimization (BO) offer an effective framework to address these challenges through latent-space Bayesian optimization (LSBO), by learning continuous latent representations of discrete peptide sequences and performing BO in this space, where smooth interpolation between sequences is possible [Gómez-Bombarelli et al., 2018]. This combination enables iterative workflows that can substantially reduce the number of experimental evaluations required to identify high affinity peptide-MHC binders. Gaussian process (GP) priors play a central role in Bayesian optimization, as they encode assumptions about the smoothness and structure of the objective function. Commonly, these priors are fitted directly on the target task, which may result in poor early-stage performance when data are scarce or domain knowledge is limited. Recent work such as HyperBO [Wang et al., 2024] proposes learning data-driven GP priors from related tasks before optimization begins. In this setting, tasks are treated as independent draws from a shared underlying process, and pre-training estimates hyperparameters that capture common structure across tasks, yielding a more informative prior for new optimization problems.

While MHC molecules are among the most polymorphic proteins in the human genome, with over 20,000 recognized classical human MHC class I alleles [Aflalo and Boyle, 2021], experimental binding affinity data exists for fewer than 300 [Vita et al., 2025]. In the absence of data for a specific allele, existing LSBO methods commonly initialize optimization using randomly generated samples, leading to inefficient early exploration. There is currently no method to effectively exploit informative binding affinity data from related alleles in existing approaches, thereby underutilizing available experimental knowledge.

In this work, we propose PepCABO (Peptide Contrastive-Aligned Bayesian Optimization), an LSBO framework for peptide-MHC affinity optimization that explicitly addresses the limitations of existing LSBO methods. PepCABO is built on a dual variational autoencoder (dual-VAE) architecture that jointly models peptide and MHC allele representations, implements knowledge transfer across alleles, and provides sample efficient optimization. The proposed training method promotes the latent representations of alleles to align with the latent representations of its strong peptide binders, enabling structured information sharing across alleles. The pretrained latent space and surrogate guide early stages of Bayesian optimization toward regions with high objective values, improving sample efficiency under limited budgets. We evaluate PepCABO under both high and low experimental budget settings against baseline methods and demonstrate state-of-the-art performance in both settings.

## 2. Material and Methods

### 2.1. Background

#### 2.1.1. Latent-space Bayesian Optimization

Bayesian optimization (BO) is a widely used framework for optimizing expensive black-box functions by iteratively fitting a probabilistic surrogate model, mainly Gaussian processes (GP), and selecting new evaluation points using an acquisition function [Frazier, 2018, Wang et al., 2023]. Despite its effectiveness, direct application of BO to discrete domains such as biological sequences is challenging due to the combinatorial size of the search space and the absence of a smooth structure.

Latent-space Bayesian optimization (LSBO) addresses this limitation by learning a continuous, low-dimensional latent representation **z** ∈*Ƶ* = ℝ^*d*^ of discrete objects **x** ∈ 𝒳 (here peptide sequences) using generative models, most commonly variational autoencoders (VAEs), and performing BO in the resulting latent space [Gómez-Bombarelli et al., 2018]. Formally, VAE is a neural architecture consisting of a deep generative latent variable model *p*_*θ*_(**x, z**) = *p*_*θ*_(**x**|**z**)*p*(**z**) that is trained using amortized variational inference with an encoder model *q*_*ϕ*_(**z**|**x**). The encoder *q*_*ϕ*_(**z**|**x**) and the decoder *p*_*θ*_(**x**|**z**) map between discrete sequences **x** and continuous latent variables **z**. Having the dataset 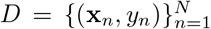 of sequences **x**_*n*_ and their objective score *y*_*n*_, a surrogate model is trained on the latent dataset 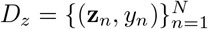, where 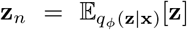 to approximate the unknown objective function *f* : 𝒳 → ℝ as a function of the latent variables, such that *f* (**x**_*n*_) ≈ *g*(**z**_*n*_). An acquisition function is used to propose a batch of promising latent candidates 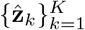 by maximizing an acquisition criteria computed from the surrogate posterior, balancing exploitation and exploration. The latent representations are mapped back to the input space by 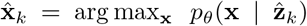, and then evaluated using the expensive objective function, yielding 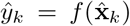. Following the evaluation, the latent dataset is updated by incorporating the newly evaluated pairs, 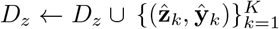. Next, the surrogate model is refitted on the updated dataset, and this procedure is repeated until a termination criterion is met.

#### 2.1.2. Joint Representation Learning for LSBO

Classical LSBO methods train the generative model and the surrogate model independently, resulting in a latent space organized purely for reconstruction without alignment to the black-box objective function. This mismatch can lead to inaccurate surrogate modeling, since similarity in latent space does not necessarily imply similarity in objective space. *Correlated latent space Bayesian Optimization (CoBO)* [Lee et al., 2023] introduces an additional step for LSBO that jointly optimizes the VAE and the surrogate model after *n* consecutive optimization steps without improvement in the best observed value. The aim is to enforce a strong correlation between distances in the latent space and differences in objective values, thereby inducing a latent geometry that is more compatible with Bayesian optimization.

To achieve this, *CoBO* introduces a new loss ℒ, in which the surrogate objective is augmented with the standard VAE loss and additional regularization terms that incentivize Lipschitz continuity of the objective with respect to the latent variables:

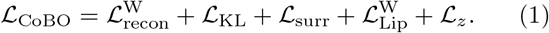

Here, 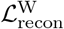 denotes the weighted reconstruction loss, which prioritizes accurate reconstruction of high-performing samples, ℒ_KL_ is the standard KL divergence regularization, and ℒ_surr_ is the GP surrogate model regression loss. The term 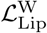 constrains the ratio between objective differences and latent distances to remain below a Lipschitz constant, with additional weighting that ensures greater smoothness in regions associated with higher objective values. The term ℒ_*z*_ regularizes pairwise latent distances to stabilize the latent space.

Building upon this, Inversion-based Latent Bayesian Optimization *(InvBO)* [Chu et al., 2024] addresses another critical limitation of LSBO methods, namely the latent input misalignment problem, which arises from imperfect reconstruction in VAE-based generative models. In this setting, a latent representation **z** associated with an observed objective value *y* = *f* (**x**) may decode to an input 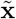 such that 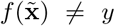. This discrepancy can degrade surrogate accuracy and acquisition reliability. *InvBO* extends *CoBO* by introducing a latent inversion procedure that performs gradient-based optimization over each triplets (**x, z**, *y*) to minimize the reconstruction loss between decoded sample **z** and previously evaluated input **x**, thereby constructing consistent triplets (**x, z**^⋆^, *y*), where **z**^⋆^ = arg max_**z**_ log *p*_*θ*_(**x** | **z**) and *f* (**x**) = *y*.

### 2.2. Proposed Method

We formulate the peptide-MHC binding affinity optimization as a black-box optimization problem over the discrete space of peptide sequences for a given target MHC allele. Let 𝒳_*p*_ denote the space of valid peptide sequences of varying lengths. Also, let 𝒳_*m*_ denote the space of MHC allele sequences or representations. Given a target allele **x**_*m*_ ∈𝒳_*m*_, our objective is to find peptide sequences that maximize the binding affinity function *f* : 𝒳_*p*_ *×* 𝒳_*m*_ → *ℝ*, where *f* (**x**_*p*_, **x**_*m*_) represents the binding affinity between peptide **x**_*p*_ and allele **x**_*m*_. In this study, to emulate *in vitro* experimental binding assays, we consider the *MHCflurry 2*.*0* predictor as the expensive black-box objective function.

#### 2.2.1. Sequence VAE

In order to introduce cross-allele information into the latent geometry and enable effective optimization, we propose a dual variational autoencoder (dual-VAE) architecture that learns coupled continuous representations of both peptides and MHC alleles via contrastive alignment. Following the LSBO methodoloy (Section 2.1.1), first a peptide VAE is defined using an encoder 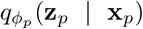 and a decoder 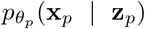 that map between discrete peptide sequences **x**_*p*_ ∈ *𝒳*_*p*_ and continuous latent representations 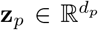. Given the limited variability in the peptide lengths **x**_*p*_, the decoder is implemented using a CNN, while a transformer encoder is used as the encoder to capture the biological nuances of the input.Similar to the peptide-VAE, an additional MHC-VAE is defined by 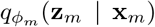 and 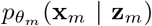, where 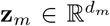 and *d*_*p*_ = *d*_*m*_. Binding is primarily determined by 34 binding-groove residues (the pseudosequence) [Nielsen et al., 2007]; we embed the full sequence with *ProtBERT* [Heinzinger et al., 2024] *and use the corresponding 34 residue embeddings as the allele representation (see Supplementary Section S1 for architectural details)*.

#### 2.2.2. Multimodal Ranked Contrastive Alignment

In order to align the MHC allele with the latent representations of its corresponding high objective value peptides, we introduce an extended multimodal contrastive loss. *Rank-N-Contrast* (RNC) [Zha et al., 2023] is a novel contrastive loss that reformulates InfoNCE by contrasting samples according to their relative rankings in the target space rather than using hard binary labels. Building on this idea, we adopt a multimodal extension, referred to as *Multi-RNC*, to align peptide and MHC latent representations.

Let *𝒳*_*m*_ denote the set of unique MHC alleles and let *N* be the number of observed triplets (**x**_*p*_, **x**_*p*_, *y*_*p,m*_) in the dataset **D** where *y*_*p,m*_ represent the objective value (i.e., binding affinity) between **x**_*p*_ and **x**_*m*_. For each allele **x**_*m*_, we denote by *𝒫*(**x**_*m*_) := { **x**_*p*_ | (**x**_*p*_, **x**_*m*_, *y*_*p,m*_) ∈**D**} the set of peptides that their corresponding objective value with the given MHC allele is available. Given an anchor pair (**x**_*p*_, **x**_*m*_), the corresponding contrastive set is

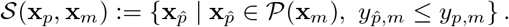

In addition, we define similarity between a peptide **x**_*p*_ and an MHC allele **x**_*m*_ by the negative Euclidean distance in the embedding space sim(**x**_*p*_, **x**_*m*_) = ∥**z**_*p*_ −**z**_*m*_∥_2_ . then the resulting Multi-RNC loss takes the form::

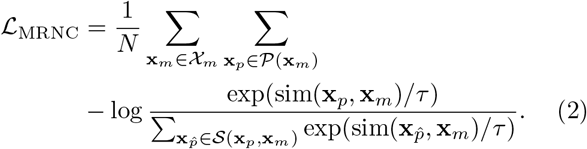

where *τ* is the temperature parameter. *ℒ*_MRNC_ encourages peptide embeddings with higher objective values to lie closer to the corresponding MHC allele than those with lower affinity, allowing a latent space geometry that reflects relative objective values and improves the efficiency of downstream Bayesian optimization.

#### 2.2.3. Surrogate Pre-training

To leverage the learned continuous latent space for efficient optimization, we train a probabilistic surrogate model to approximate the binding affinity function in the joint embedding space. Unlike *HyperBO* [Wang et al., 2024], which models each task (i.e., MHC allele **x**_*m*_ binding prediction in this setting) as an independent function under a shared prior, we define a single GP over the joint latent space (**z**_*p*_, **z**_*m*_) to share information across related alleles through the kernel structure:

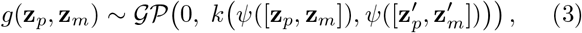

where *k*(·, · ) is a kernel function (RBF kernel in our case) and *ψ*(·) is a multilayer perceptron (MLP) operator. Since the existing datasets are very large, we adopt a sparse variational Gaussian process to model *g* in order to remain computationally scalable.

Before performing LSBO for a given MHC allele, we first train the peptide VAE and the allele VAE separately using all available peptide and allele sequences, regardless of the availability of binding affinity data, with the standard VAE loss. The peptide and allele VAEs are then pre-trained together with the surrogate GP model on triplets (**x**_*p*_, **x**_*m*_, *y*_*p,m*_) ∈ **D** using the following objective (see Supplementary Section S2.1 for detailed expressions):

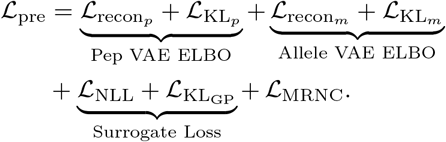

The first two terms correspond to the ELBO objectives of the peptide and allele VAEs, each consisting of a reconstruction term and a KL regularization term. The surrogate loss includes the negative log-likelihood and a KL term for the variational inducing distribution, together forming the variational evidence lower bound (ELBO) of the sparse Gaussian process. Finally, ℒ_MRNC_ aligns peptide and allele embeddings according to their relative binding affinities.

By training the peptide and allele VAEs simultaneously with the surrogate loss using ℒ_pre_ before the actual BO, the resulting geometry and surrogate model capture structured relationships across related alleles. The pretrained surrogate is then used during the subsequent Bayesian optimization stage, allowing information from previously observed alleles to inform predictions for a new target allele. This enables knowledge transfer across related alleles, improves sample efficiency, and guides exploration toward promising peptide regions.

In the context of Bayesian optimization, prediction accuracy for high binders and effective modeling of high-affinity samples are most relevant. Therefore, we introduce a data-dependent weighting function *w*(*y*_*p,m*_) (detailed definition Supplementary Section S2.2) that assigns larger weights to triplets (**x**_*p*_, **x**_*m*_, *y*_*p,m*_) with higher objective value *y*_*p,m*_ in both the negative log-likelihood 𝓁_NLL_ and the multimodal ranking loss 𝓁_MRNC_, thereby improving the local structure around MHC alleles and

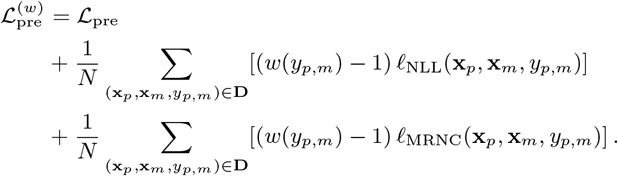

#### 2.2.4. Sample Initialization

To obtain an informative initial pool already at the first BO step, we sample peptide candidates in the learned latent space by employing the acquisition function under the pretrained surrogate model, and subsequently decode them using the peptide VAE decoder. Trust region based acquisition functions [Eriksson et al., 2019] operate by sampling around evaluated promising candidates using the surrogate to identify the next candidates for evaluation in Bayesian optimization.

Expanding on this, based on the alignment induced by *L*_MRNC_ (Eq 2) and the similarity definition in Sec 2.2.2, the latent representation of the MHC allele, 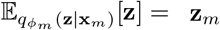, is expected to lie near high affinity peptides (to that specific MHC allele) in the peptide latent space. Therefore, rather than relying on randomly initialized samples, we center a trust region around **z**_*m*_ and sample via Thompson sampling from its neighborhood in the peptide latent space using the pretrained surrogate. This approach biases posterior sampling toward regions that are structurally close to peptides with high objective values. By drawing multiple Thompson samples from the surrogate posterior centered around **z**_*m*_, the initialization phase is guided toward promising regions of the search space, improving early stage sample efficiency.

#### 2.2.5. Bayesian Optimization Step

During Bayesian optimization for the given allele 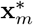, we perform end-to-end updates of the latent representations (as described in Section 2.1.2) by optimizing a combined objective consisting of the CoBO loss and the Multi-RNC alignment term. Since only one MHC allele is considered during optimization, we fix 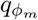 and 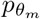, and only the peptide VAE remains learnable during BO:

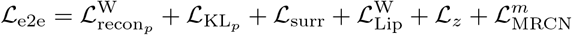

Here, 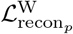 and 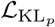 denote the weighted reconstruction loss and KL divergence of the peptide VAE, respectively. ℒ_surr_ denotes the GP surrogate loss, while 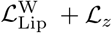 correspond to the regularization terms described previously. Finally, 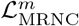 represents the ranked contrastive loss computed only on peptide-allele pairs observed during optimization (i.e., for the single target allele **x**^*\**^). A detailed description of all loss terms and weighting schemes is provided in Supplementary Section S2.3.

In order to update the surrogate after evaluating new candidates, during the optmization, the pretrained surrogate model is updated while preserving the informative prior learned during pre-training as much as possible. To achieve this, we freeze the kernel hyperparameters, likelihood noise, and feature extractor, and update only the variational posterior parameters and inducing point locations.

### 2.3. Dataset

All peptide-MHC pairs used during the dual-VAE training were extracted from the *MHCflurry 2*.*0* training dataset [O’Donnell et al., 2020]. This dataset contains 162 unique MHC class I alleles and roughly 600,000 unique peptide-allele pairs. The training data include both quantitative affinity measurements (IC50 values) and qualitative monoallelic mass-spectrometry (MS) ligand data, of which we focus on binding affinity. For the purposes of evaluating BO methods, we need to be able to evaluate the black-box objective function *f* (**x**_*p*_, **x**_*m*_) for *any* input pair **x**_*p*_ ∈*𝒳*_*p*_ and **x**_*m*_ ∈*𝒳*_*m*_. Therefore, we defined the black-box objective function using *MHCflurry 2*.*0*, as described in Section 2.4.1. Using a predictive model as a proxy of the binding affinity enables fully *in silico* emulation of BO experiments, allowing systematic evaluation of optimization strategies without requiring wet-lab assays at each iteration.

Approximately 40% of all pairs belong to only 12 specific MHC alleles, resulting in a substantial data imbalance. The distribution of allele frequencies and the names of the 12 held-out alleles are provided in the Supplementary Section S3. During pretraining and BO emulations, we treated these 12 alleles as if no binding affinity data were available for them, and the surrogate was pretrained exclusively on the remaining 143 alleles. The peptide optimization using BO is then done for the left-out 12 alleles. This framework ensures that, during BO emulation, the BA predictor maintains maximal accuracy, since it was originally trained on these alleles with a substantial amount of data. Moreover, this abundance of experimental data provides a well-characterized baseline against which we can compare our in silico optimization results, and finally, it prevents the surrogate from overfitting to these specific alleles.

### 2.4. Objective Function and Evaluation Metrics

#### 2.4.1 MHCflurry

MHCflurry 2.0 is an established pan-allele predictor for MHC class I peptide binding and presentation [O’Donnell et al., 2020], which provides two main outputs for each peptide-MHC pair: a binding affinity (BA) prediction and a presentation score (PS). The binding affinity predictor estimates the peptide-MHC interaction strength using a neural network ensemble trained on experimental affinity measurements and monoallelic mass spectrometry data. The BA predictor networks are trained on a log-transformed scale, given by *s*_BA_ = 1 − log_50,000_(IC50), where IC50 denotes the binding affinity in nanomolar (nM). Higher values of *s*_BA_ indicate stronger predicted binding.

The presentation score (PS) integrates binding affinity with antigen processing effects by combining the binding affinity (BA) prediction with an antigen processing (AP) predictor using a logistic regression model trained on multiallelic mass spectrometry datasets, producing a probability-like score in [0, 1] that reflects the likelihood of a peptide being presented on the cell surface. We primarily focus on binding affinity as the optimization objective rather than the presentation score, since it is a physically meaningful and experimentally measurable quantity *in vitro*. This choice enables direct translation of PepCABO to real world experimental settings, where Bayesian optimization can be performed using laboratory measurements instead of relying on proxy scores that exist only within predictive models.

To ensure a consistent and continuous optimization target, the *MHCflurry 2*.*0* predictor was applied to all peptide-allele pairs in the training dataset, regardless of data type, to obtain unified objective values (i.e., binding affinity (BA) or presentation score (PS)). Consequently, the training dataset was represented as triplets of the form (**x**_*p*_, **x**_*m*_, *y*_*p,m*_), where *y*_*p,m*_ = *f* (**x**_*p*_, **x**_*m*_) denotes the objective evaluated by *MHCflurry 2*.*0*.

#### 2.4.2. Evaluation Metrics

To compare the performance of different models across varying data regimes and objectives, we report the area under the best-so-far objective curve across oracle evaluations (AUOC). AUOC captures both sample efficiency and convergence behavior, reflecting how rapidly and consistently a method improves under a fixed evaluation budget. We also report the best objective value found within each budget, as well as the best value observed at the midpoint of each optimization trajectories, providing additional insight of performance within each budget.

## 3. Experiments

### 3.1. Baselines

We compare PepCABO against vanilla LSBO and InvBO, described in Section 2.1, implemented for each MHC allele independently using only the peptide-VAE with the same trust-region-based search, without any pretraining, and with a surrogate model as in Eq. 3 using only *z*_*p*_ as input, as well as against PepPPO [Chen et al., 2023]. While LSBO and InvBO operate in a learned latent space using Bayesian optimization, PepPPO is a reinforcement learning (RL) framework for de novo generation of MHC class I binding peptides via iterative sequence mutation. Starting from a random peptide and a target MHC allele, an agent mutates one amino acid at each step and receives a reward only at terminal states based on the predicted PS of the resulting peptide using MHCflurry 2.0.

PepPPO’s policy is optimized through iterative interaction with the mutation environment, during which the agent learns a mutation policy that transforms random peptides into sequences predicted to have high objective value. As a result, PepPPO relies on a large number of oracle evaluations during training. Each mutation step produces a new peptide-MHC pair that is typically absent from the training dataset and therefore requires a new query to obtain a reward signal. Due to the long mutation trajectories and the large number of episodes used during training, this approach requires up to 10,000,000 oracle calls across all alleles supported by MHCflurry 2.0 (approximately 10,000 alleles) prior to evaluation. In contrast, neither our method nor other BO baselines requires oracle calls before optimization and instead relies solely on existing data from 155 alleles for pre-training. Consequently, while PepPPO is practical when the oracle is a fast *in silico* predictor such as MHCflurry, it is not well suited for scenarios involving expensive experimental measurements, such as *in vitro* binding affinity assays.

### 3.2. Results

The PepCABO is compatible with any LSBO framework. In this study, we implemented it within the InvBO framework, selected based on its strong performance in prior benchmark studies. All methods were trained, and evaluated under both the log-transformed binding affinity (BA) and presentation score (PS) objectives. While PepPPO was originally trained on the PS objective, we retrained it on BA as well to enable a fair comparison across both settings. For PepCABO, we conducted experiments with both random initialization (i.e., by randomly selecting from the set of all existing peptides in the pretraining dataset) and guided initialization, to assess performance independently of the starting point.

All model evaluations were performed under two budget settings to reflect both realistically constrained and less constrained experimental scenarios. The low-budget setting allowed a total of 200 oracle calls, while the high-budget setting allowed 1,000 oracle calls. Full details of the budget configuration for each method are provided in the Supplementary Section S4.

Results for both objective values and both budget regimes are summarized in Table 1, with representative optimization trajectories for the binding affinity objective shown in Fig. 1. PepCABO consistently shows improved performance compared to the baseline methods, regardless of the initialization strategy and budget. However, with guided initialization, it is evident that Bayesian optimization converges much faster and reaches better solutions within fewer iterations. For instance, PepCABO’s guided initialization in the low-budget setting finds a better solution in its second batch compared to the baselines overall.In Supplementary Section S5, we present an ablation study showing that each component of PepCABO contributes to the overall performance.

**Table 1:**
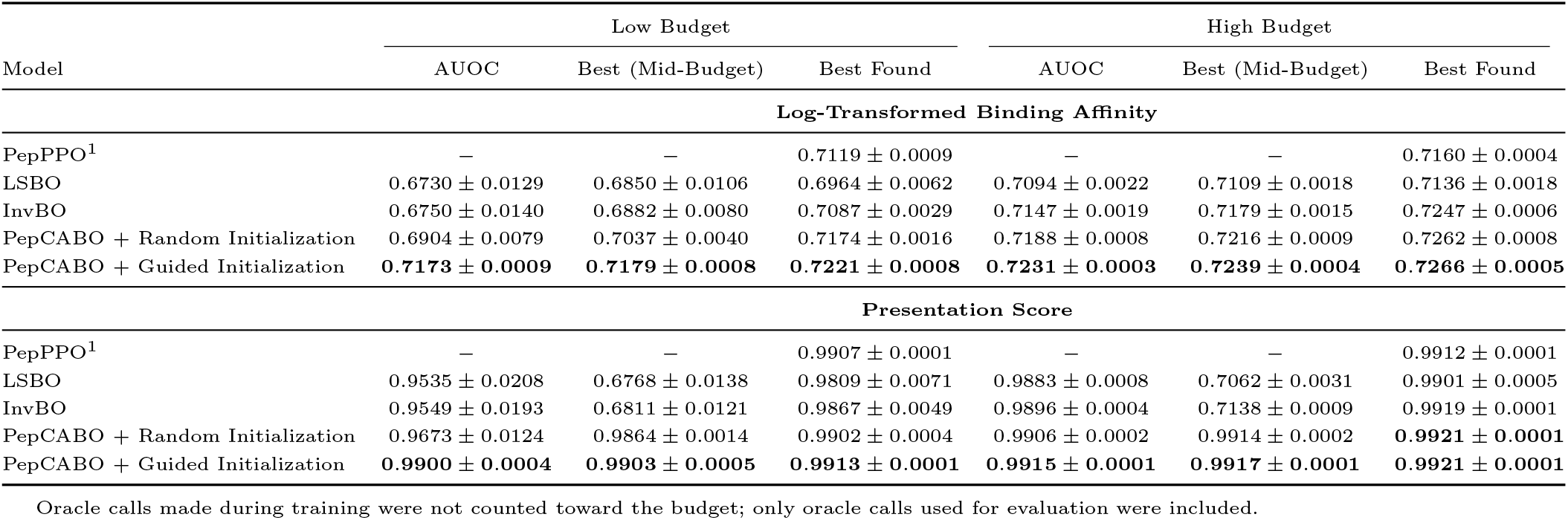
Summary of experimental results under different budget settings. All experiments are performed independently for each of the 12 held-out alleles and repeated across 10 random seeds. For each seed, performance is first averaged across alleles; the reported mean and standard deviation are then computed across seeds.

**Figure 1:**
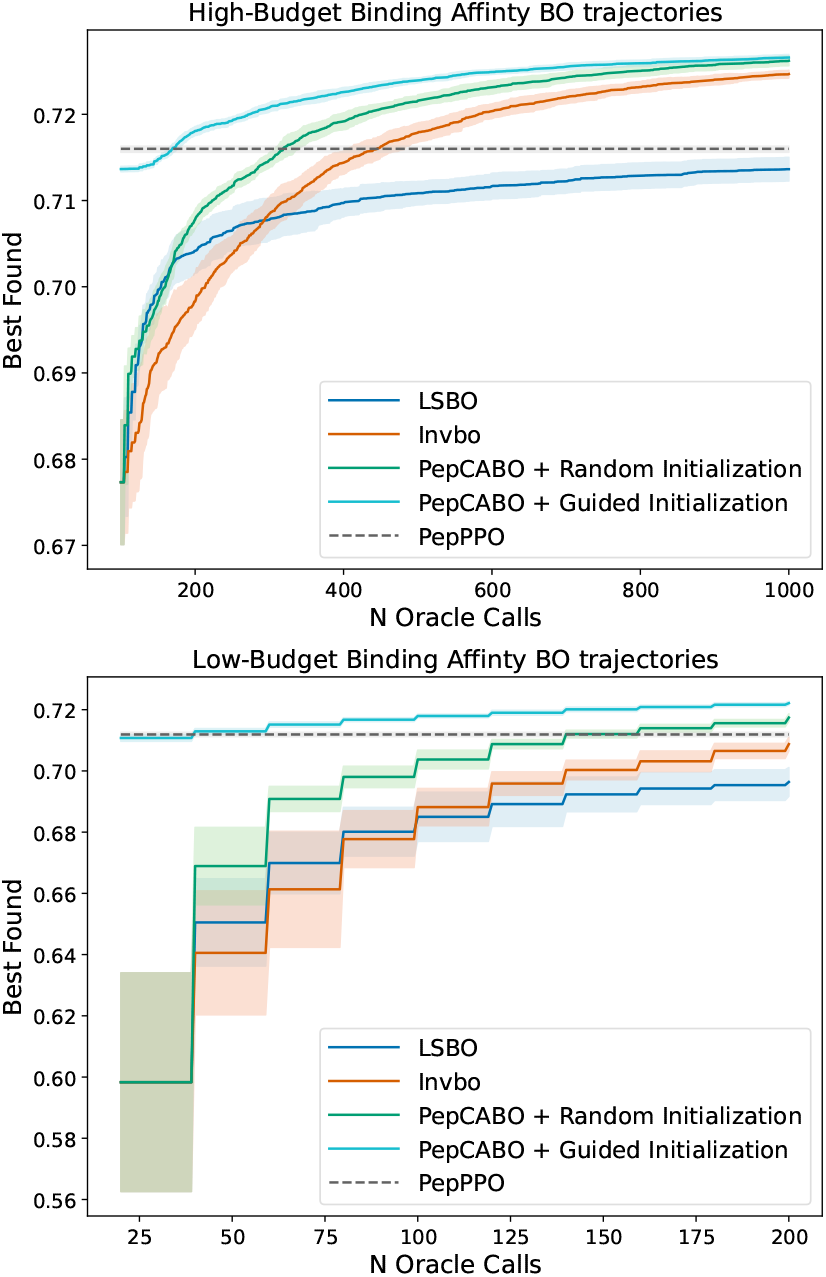
Bayesian optimization trajectories in the high-budget (up) and low-budget (down) settings, showing the best observed log-transformed BA as a function of the number of oracle calls. Shaded regions indicate 95% confidence intervals. The PepPPO line indicates the best value observed after spending the designated budget; it is not shown as a function of oracle calls and is instead presented as a constant reference line.

### 3.3. Initialization Evaluation

Experimental validation of the PepCABO through *in vitro* assays is beyond the scope of this study. Nevertheless, PepCABO’s effectiveness can be assessed indirectly by evaluating the impact of different initialization strategies, since better initialization typically leads to faster convergence. For this analysis, we use the same proposed training objective described in Section 2.2.3. However, in this setting, the surrogate model is trained using only the quantitative portion of the data, and a modified MRNC loss is employed that is able to handle both qualitative and quantitative signals when available. Specifically, we compare two approaches: random initialization and guided initialization following our methodology using experimental data. (details of modified loss and random initialization method for experimental values are provided in the Supplementary Section S6 and Section S7)

Since experimental *IC*_50_ values may follow a different distribution than predicted *IC*_50_ estimates, we assess the initialization batch using percentile ranks instead of raw values. Specifically, for each MHC allele **x**_*m*_, we compute the percentile rank of both the average objective value of the initialization batch and the best observed sample relative to a reference set of peptides. In the experimental setting, percentile ranks are computed with respect to peptides with quantitative measurements available for the corresponding allele, whereas in the emulation setting they are computed relative to MHCflurry-predicted values for the same reference set used in the experimental setting.

As demonstrated in Table 2, guided initialization consistently produces a substantially more informative initial batch than random sampling. Under MHCflurry-predicted values, the batch average reaches the 88.5th percentile of the reference distribution, compared to only the 23.5th percentile for random initialization, while the batch maximum reaches the 99.8th percentile, compared to 79.1th under random sampling. A similar pattern is observed when evaluated quantitatively against experimental measurements. The guided initialization achieves a batch average at the 73.0th percentile, compared to 49.9th for random initialization, while the batch maximum reaches the 97.9rd percentile, compared to 95.2st for randomly selected batches.

**Table 2:** Initialization performance using experimental or MHCflurry-predicted values. Metrics are percentile ranks (left) and log-*IC*_50_ (right).

| Init | Percentile |  | Log-IC<sub>50</sub> |  |
| --- | --- | --- | --- | --- |
|  | Avg | Max | Avg | Max |
| **Experimental** |  |  |  |  |
| Random | 49.9±1.7 | 95.2±2.0 | 0.39±0.02 | 0.91±0.05 |
| Guided | 73.0±1.3 | 97.9±0.5 | 0.63±0.02 | 1.12±0.09 |
| **MHCflurry** |  |  |  |  |
| Random | 23.5±1.2 | 79.1±8.3 | 0.19±0.01 | 0.60±0.05 |
| Guided | 88.5±1.4 | 99.8±0.1 | 0.65±0.01 | 0.71±0.00 |

## 4. Discussion

We introduced a dual-VAE latent-space Bayesian optimization model that leverages biological cross allele structure to improve sample efficiency in peptide-MHC affinity optimization. Across 12 common held out alleles, PepCABO consistently outperformed LSBO and InvBO as well as the RL baseline (PepPPO) in both the low and high budget regimes, achieving stronger best-found binding affinity and faster convergence. Notably, based on available experimental measurements, we demonstrated that the performance improvement provided by the proposed initialization strategy is comparable to that observed in the emulated scenario, indicating that the method is compatible with experimental settings and is not limited to *in silico* predictors. A key property of peptide-MHC datasets is that they consisted of *quantitative* affinity measurements (e.g., IC_50_) and *qualitative* ligand evidence from mass spectrometry. In this work, we used MHCflurry predictions as a unified surrogate objective to enable systematic and reproducible evaluation of BO methods. However, when training directly on experimental data, using a *censored surrogate* that explicitly models qualitative observations as bounded, while treating quantitative IC_50_ values as exact targets, could make a significant improvement. Such a methodology would better utilize mixed experimental data and would be able to enhance accuracy and sample efficiency.

## Author Contributions

M.G., D.K., A.D., and H.L. designed the study. M.G. developed the methodology, implemented the models, conducted experiments, analyzed the results, and wrote the first draft. D.K. and A.D. provided methodological feedback and contributed to manuscript revisions. H.L. supervised the project and provided conceptual guidance and manuscript feedback.

## Acknowledgments

We acknowledge the computational resources provided by the Aalto Science-IT project. This work is supported in part by funds from the Research Council of Finland (#359135), the Cancer Foundation of Finland, and the Sigrid Jusélius Foundation.

## References

A. Aflalo and L. H. Boyle. Polymorphisms in mhc class i molecules influence their interactions with components of the antigen processing and presentation pathway. International Journal of Immunogenetics, 48(4):317–325, 2021. doi: 10.1111/iji.12546.

Z. Chen, B. Zhang, H. Guo, P. Emani, T. Clancy, C. Jiang, M. Gerstein, X. Ning, C. Cheng, and M. R. Min. Binding peptide generation for mhc class i proteins with deep reinforcement learning. Bioinformatics, 39(2):btad055, 01 2023. ISSN 1367-4811. doi: 10.1093/bioinformatics/btad055.

J. Chu, J. Park, S. Lee, and H. J. Kim. Inversion-based latent bayesian optimization. In Proceedings of the 38th International Conference on Neural Information Processing Systems, NIPS ‘24, Red Hook, NY, USA, 2024. Curran Associates Inc. ISBN 9798331314385.

L. Collesano, M. Luksza, and M. Lässig. Energy landscapes of peptide-mhc binding. PLOS Computational Biology, 20(9):1–19, 09 2024. doi: 10.1371/journal.pcbi.1012380.

J. Comber and R. Philip. MHC class I antigen presentation and implications for developing a new generation of therapeutic vaccines. Ther Adv Vaccines, 2(3):77–89, May 2014.

D. Eriksson, M. Pearce, J. R. Gardner, R. Turner, and M. Poloczek. Scalable global optimization via local Bayesian optimization. Curran Associates Inc., Red Hook, NY, USA, 2019.

P. I. Frazier. Bayesian Optimization, chapter 11, pages 255–278. INFORMS, 2018. doi: 10.1287/educ.2018.0188.

R. Gómez-Bombarelli, J. N. Wei, D. Duvenaud, J. M. Herńandez-Lobato, B. Sanchez-Lengeling, D. Sheberla, J. Aguilera-Iparraguirre, T. D. Hirzel, R. P. Adams, and A. Aspuru-Guzik. Automatic chemical design using a data-driven continuous representation of molecules. ACS Central Science, 4(2):268–276, Feb 2018. ISSN 2374-7943. doi: 10.1021/acscentsci.7b00572.

A. K. Haj, M. E. Breitbach, D. A. Baker, M. S. Mohns, G. K. Moreno, N. A. Wilson, V. Lyamichev, J. Patel, K. L. Weisgrau, D. M. Dudley, and D. H. O’Connor. High-throughput identification of MHC class I binding peptides using an ultradense peptide array. J. Immunol., 204(6):1689–1696, Mar. 2020.

M. Heinzinger, K. Weissenow, J. G. Sanchez, A. Henkel, M. Mirdita, M. Steinegger, and B. Rost. Bilingual language model for protein sequence and structure. NAR Genomics and Bioinformatics, 6(4):lqae150, 11 2024. ISSN 2631-9268. doi: 10.1093/nargab/lqae150.

S. Lee, J. Chu, S. Kim, J. Ko, and H. J. Kim. Advancing bayesian optimization via learning correlated latent space. In Proceedings of the 37th International Conference on Neural Information Processing Systems, NIPS ‘23, Red Hook, NY, USA, 2023. Curran Associates Inc.

M. Nielsen, C. Lundegaard, T. Blicher, K. Lamberth, M. Harndahl, S. Justesen, G. Røder, B. Peters, A. Sette, O. Lund, and S. Buus. NetMHCpan, a method for quantitative predictions of peptide binding to any HLA-A and -B locus protein of known sequence. PLoS One, 2(8):e796, Aug. 2007.

T. J. O’Donnell, A. Rubinsteyn, and U. Laserson. Mhcflurry 2.0: Improved pan-allele prediction of mhc class i-presented peptides by incorporating antigen processing. Cell Systems, 11(1):42–48.e7, 2020. ISSN 2405-4712. doi: 10.1016/j.cels.2020.06.010.

R. Vita, N. Blazeska, D. Marrama, S. Duesing, J. Bennett, J. Greenbaum, M. De Almeida Mendes, J. Mahita, D. K. Wheeler, J. R. Cantrell, J. A. Overton, D. A. Natale, A. Sette, and B. Peters. The immune epitope database (iedb): 2024 update. Nucleic Acids Research, 53(D1): D436–D443, Jan. 2025. doi: 10.1093/nar/gkae1092.

X. Wang, Y. Jin, S. Schmitt, and M. Olhofer. Recent advances in bayesian optimization. ACM Comput. Surv., 55(13s), July 2023. ISSN 0360-0300. doi: 10.1145/3582078.

Z. Wang, G. E. Dahl, K. Swersky, C. Lee, Z. Nado, J. Gilmer, J. Snoek, and Z. Ghahramani. Pre-trained gaussian processes for bayesian optimization. J. Mach. Learn. Res., 25(1), Jan. 2024. ISSN 1532-4435.

M. Wieczorek, E. T. Abualrous, J. Sticht, M. Álvaro Benito, S. Stolzenberg, F. Nóe, and C. Freund. Major histocompatibility complex (mhc) class i and mhc class ii proteins: Conformational plasticity in antigen presentation. Frontiers in Immunology, Volume 8 - 2017, 2017. ISSN 1664-3224. doi: 10.3389/fimmu.2017.00292.

K. Zha, P. Cao, J. Son, Y. Yang, and D. Katabi. Rank-n-contrast: Learning continuous representations for regression. In Thirty-seventh Conference on Neural Information Processing Systems, 2023.

