## Supplementary File for "PepCABO: Latent-space Bayesian Optimization for Peptide-MHC Binding Using Contrastive Alignment"

#### 1 Neural Network Architectures

##### 1.1 Peptide Variational Autoencoder

**Encoder.** Peptide sequences are padded to a fixed length and embedded using a fixed amino acid substitution matrix (BLOSUM62 embedding), followed by projection to a hidden dimension of 128. The projected representations are processed by a positional encoding layer and a transformer encoder consisting of 6 layers with 8 attention heads, feedforward dimension 256, and dropout rate 0.1. Transformer token representations are aggregated via masked average pooling to obtain sequence embedding, which is mapped to the mean $\mu(x)$ and standard deviation $\sigma(x)$ of a 64-dimensional diagonal Gaussian. Latent variables are sampled using the reparameterization trick.

**Decoder.** The decoder reconstructs peptide sequences from the latent vector using a convolutional architecture. The latent representation is first linearly projected to a fixed-length feature map and then processed by three one-dimensional transposed convolutional layers with channel progression $24 \rightarrow 64 \rightarrow 32 \rightarrow$ vocabulary dimension. ReLU activations, dropout (rate 0.15), and layer normalization are applied between layers. The final output consists of position-wise logits over the amino acid vocabulary for peptides up to length 15, optimized using cross-entropy loss.

##### 1.2 Allele Variational Autoencoder

**Encoder.** MHC alleles are represented by their pseudosequences. The full sequence is first encoded using a pretrained protein language model, and the embeddings corresponding to the 34 pseudosequence positions are then extracted (1024 dimensions per position). The resulting $34 \times 1024$ representation is processed by a 1D CNN encoder with two convolutional blocks (kernel size 9), using channel sizes $1024 \rightarrow 64 \rightarrow 128$, with ReLU activations and dropout. The final feature map is flattened and mapped to the mean $\mu(x)$ and standard deviation $\sigma(x)$ of a 64-dimensional diagonal Gaussian latent space. Latent variables are sampled using the reparameterization trick.

**Decoder.** The decoder maps the latent vector through a fully connected layer followed by a lightweight transposed 1D CNN (channels $128 \rightarrow 64 \rightarrow$ vocabulary) to reconstruct residue pseudosequence, optimized using cross-entropy loss.

#### 2 Loss Function Details

##### 2.1 Full Expression of the Pretraining Objective

During pretraining, the peptide VAE, allele VAE, and the sparse Gaussian process (GP) surrogate are jointly optimized using the following objective:

$$\mathcal{L}_{\text{pre}} = \underbrace{\mathcal{L}_{\text{recon}_p} + \mathcal{L}_{\text{KL}_p}}_{\text{Peptide VAE ELBO}} + \underbrace{\mathcal{L}_{\text{recon}_m} + \mathcal{L}_{\text{KL}_m}}_{\text{Allele VAE ELBO}} + \underbrace{\mathcal{L}_{\text{NLL}} + \mathcal{L}_{\text{KL}_{\text{GP}}}}_{\text{Surrogate Loss}} + \mathcal{L}_{\text{MRNC}}.$$

The peptide VAE evidence lower bound (ELBO) terms are defined as

$$\mathcal{L}_{\text{recon}_p} = -\mathbb{E}_{q_{\phi_p}(z_p|x_p)}\left[\log p_{\theta_p}(x_p|z_p)\right], \qquad \mathcal{L}_{\text{KL}_p} = \text{KL}\big(q_{\phi_p}(z_p|x_p) \,\|\, \mathcal{N}(0, I)\big). \tag{1}$$

Similarly, the allele VAE ELBO terms are defined as

$$\mathcal{L}_{\text{recon}_m} = -\mathbb{E}_{q_{\phi_m}(z_m|x_m)}\left[\log p_{\theta_m}(x_m|z_m)\right], \qquad \mathcal{L}_{\text{KL}_m} = \text{KL}(q_{\phi_m}(z_m|x_m) \,\|\, \mathcal{N}(0, I)). \tag{2}$$

The surrogate loss corresponds to the negative variational ELBO of the sparse GP and consists of two terms:

$$\mathcal{L}_{\text{NLL}} = -\mathbb{E}_{q(f)}\left[\log p(y_{p,m} \mid f(z_p, z_m))\right], \qquad \mathcal{L}_{\text{KL}_{\text{GP}}} = \text{KL}(q(u) \,\|\, p(u)), \tag{3}$$

where $q(f)$ denotes the variational posterior over the latent function values induced by the variational distribution over the inducing variables $q(u)$, i.e., $q(f) = \int p(f \mid u)q(u)du$.

The multimodal ranked contrastive alignment term $\mathcal{L}_{\text{MRNC}}$ is defined in Eq. (2) of the main manuscript.

##### 2.2 Full expressions for the end-to-end BO objective

The end-to-end loss used during BO optimization is defined as

$$\mathcal{L}_{\text{e2e}} = \mathcal{L}^{\text{W}}_{\text{recon}_p} + \mathcal{L}_{\text{KL}_p} + \mathcal{L}_{\text{surr}} + \mathcal{L}^{\text{W}}_{\text{Lip}} + \mathcal{L}_z + \mathcal{L}^{m}_{\text{MRNC}}.$$

Following CoBO [Lee et al., 2023], each observed peptide $\mathbf{x}_p$ is assigned a weight based on its objective value $y_{p,m^*}$ with respect to the target allele $\mathbf{x}^*_m$, emphasizing promising regions of the search space:

$$w(y_{p,m^*}) = \mathbb{P}(Y > y_q), \quad Y \sim \mathcal{N}(y_{p,m^*}, \sigma^2), \tag{4}$$

where $y_q$ denotes a chosen quantile (95th percentile in our case), and $\sigma = 0.1$ is a smoothing hyperparameter.

During end-to-end BO updates, only the peptide VAE is optimized. Its weighted reconstruction term is

$$\mathcal{L}^{\text{W}}_{\text{recon}_p} = -w(y_{p,m^*})\mathbb{E}_{q_{\phi_p}(z_p|\mathbf{x}_p)}\left[\log p_{\theta_p}(\mathbf{x}_p|z_p)\right], \tag{5}$$

with KL regularization $\mathcal{L}_{\text{KL}_p}$ defined as in Eq. 1. The sparse GP surrogate is trained via its negative variational ELBO, using a formulation same as Eq. 3:

$$\mathcal{L}_{\text{surr}} = L_{\text{NLL}} + L_{\text{KL}_{\text{GP}}}. \tag{6}$$

To encourage alignment between latent distances and objective differences, we impose Lipschitz regularization [Lee et al., 2023]:

$$\mathcal{L}_{\text{Lip}} = \sum_{i,j} \max\left(0, \frac{|y_{p_i,m^*} - y_{p_j,m^*}|}{\|z_{p_i} - z_{p_j}\|_2} - L\right), \tag{7}$$

where $L$ is the median slope across sample pairs. To prioritize high-objective regions, we further apply weighting:

$$\mathcal{L}_{\text{Lip}}^{\text{W}} = \sum_{i,j} \sqrt{w(y_{p_i,m^*})w(y_{p_j,m^*})} \max\left(0, \frac{|y_{p_i,m^*} - y_{p_j,m^*}|}{\|z_{p_i} - z_{p_j}\|_2} - L\right). \tag{8}$$

To prevent excessive expansion or collapse of the latent space, we regularize the average pairwise distance [Lee et al., 2023]:

$$\mathcal{L}_z = \left| \frac{1}{N^2} \sum_{i,j} \|z_{p_i} - z_{p_j}\|_2 - c \right|, \tag{9}$$

where

$$c = 2\frac{\Gamma\left(\frac{k+1}{2}\right)}{\Gamma\left(\frac{k}{2}\right)}, \tag{10}$$

with $k = 64$ denoting the latent dimension.

During Bayesian optimization for a fixed target allele $\mathbf{x}_m^*$, we additionally impose a ranked contrastive alignment term to preserve the objective-aware latent geometry. Unlike $\mathcal{L}_{\text{pre}}$, this term is computed only over peptide–allele pairs observed during optimization for the target allele. Let $\mathcal{X}_m^*$ denote the set of peptides already evaluated by the expensive black-box objective. For an anchor peptide $\mathbf{x}_p \in \mathcal{X}_m^*$, the ranked contrastive set is defined as

$$\mathcal{S}(\mathbf{x}_p) = \{\mathbf{x}_{\hat{p}} \in \mathcal{X}_m^* \mid y_{\hat{p},m^*} \leq y_{p,m^*}\}. \tag{11}$$

The allele-specific Multi-Rank-N-Contrast (MRNC) loss is

$$\mathcal{L}_{\text{MRNC}}^m = -\frac{1}{N} \sum_{\mathbf{x}_p \in \mathcal{X}_m^*} \log \frac{\exp(\text{sim}(\mathbf{x}_p, \mathbf{x}_m^*)/\tau)}{\sum_{\mathbf{x}_{\hat{p}} \in \mathcal{S}(\mathbf{x}_p)} \exp(\text{sim}(\mathbf{x}_{\hat{p}}, \mathbf{x}_m^*)/\tau)}. \tag{12}$$

Here $\tau$ is a temperature parameter and $\text{sim}(\cdot,\cdot)$ is defined in the main manuscript. During BO, the allele encoder $q_{\phi_m}$ and decoder $p_{\theta_m}$ are frozen, and gradients propagate only through the peptide encoder $q_{\phi_p}$.

#### 2.3 Weighted loss formulation details during pre-training

Similar to Section 2.2, we use an objective-dependent weighting function during pretraining. Let $y_{p,m}$ denote the objective value between peptide $\mathbf{x}_p$, and MHC allele $\mathbf{x}_m$. However, unlike the adaptive threshold used during end-to-end training, the pretraining weights are defined using fixed thresholds based on the objective, which corresponds either to binding affinity (BA) or presentation score (PS).

**Binding Affinity (BA).** Binding affinity values are modeled on the log-transformed scale during training. An affinity of 100 nM is a widely adopted threshold and is commonly used as an indicator of strong MHC binding. Under our log-scale transformation, this corresponds to

$$y_{\text{BA}}^{\text{thr}} = 1 - \frac{\log 100}{\log 50000} \approx 0.574. \tag{13}$$

Therefore, the weighting scheme becomes

$$w(y_{p,m}) = \mathbb{P}(Y > y_{\text{BA}}^{\text{thr}}), \quad Y \sim \mathcal{N}(y_{p,m}, \sigma^2), \tag{14}$$

where $\sigma = 0.15$.

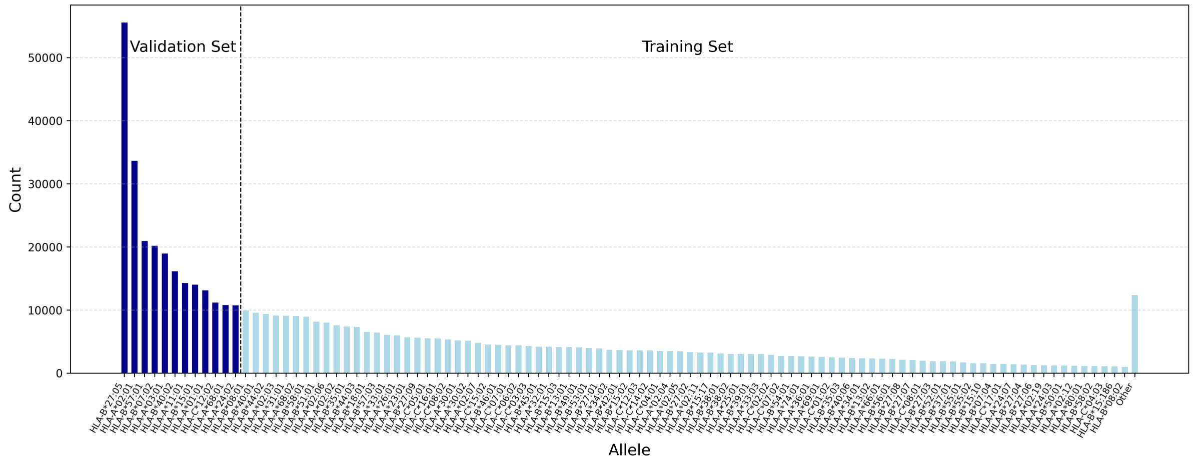


Figure 1: Distribution of peptide-allele pair counts for the top 100 most frequent MHC class I alleles in the MHCflurry 2.0 dataset.

**Presentation Score (PS).** For presentation score, which lies in $[0, 1]$, we set $y_{\mathrm{BA}}^{\mathrm{thr}} = 0.90$, determined empirically, to identify highly presented peptides. The same smoothing hyperparameter $\sigma = 0.15$ is used.

### 3 Allele Frequency Distribution

To illustrate the allele-level data imbalance in the MHCflurry 2.0 [O'Donnell et al., 2020] training dataset, the number of peptide–allele pairs available for each MHC class I allele is shown in Figure 1 for the top 100 most frequent alleles. The alleles are sorted in descending order according to the number of associated peptide–allele pairs. The dashed vertical line separates the 12 most frequent alleles (left) from the remaining alleles (right).

The full list of allele names corresponding to the top 12 most frequent alleles, which were used during the evaluation of the proposed method and the baselines, is provided in Table 1.

Table 1: Top 12 most frequent MHC class I alleles in the MHCflurry 2.0 dataset.

| | | | |
|---|---|---|---|
| HLA-B*27:05 | HLA-A*02:01 | HLA-B*57:01 | HLA-B*07:02 |
| HLA-A*03:01 | HLA-B*40:02 | HLA-A*11:01 | HLA-B*15:01 |
| HLA-A*01:01 | HLA-C*12:02 | HLA-A*68:01 | HLA-A*24:02 |

### 4 Budget Configuration Details

**High-Budget Setting.** In the high-budget regime, the total number of allowed oracle calls was set to 1,000. Of these, 100 peptides were used for initialization, and subsequent optimization proceeded in batches of size 5. After 10 consecutive BO steps without any improvement in the best-performing peptide (i.e., the peptide with the highest objective value), an end-to-end training step was applied. In this setting, no strict constraint was imposed on the number of newly evaluated peptides per step; due to the generation of previously evaluated candidates, the number of unique peptides evaluated per iteration could be fewer than 5.

**Low-Budget Setting.** To emulate more realistic experimental conditions, the low-budget regime limited each Bayesian optimization run to a total of 200 oracle evaluations. This consisted of an initial set of 20 peptides, followed by 9 optimization steps, with exactly 20 newly

Table 2: Ablation study under low- (200 calls) and high-budget (1,000 calls) settings, reporting log-transformed binding affinity (BA). Results are averaged across 12 alleles and 10 random seeds (mean ± std).

| Model | Low Budget | | | High Budget | | |
|---|---|---|---|---|---|---|
| | AUOC | Mid | Best | AUOC | Mid | Best |
| **Random Initialization** | | | | | | |
| InvBO | $0.6750 \pm 0.0140$ | $0.6882 \pm 0.0080$ | $0.7087 \pm 0.0029$ | $0.7147 \pm 0.0019$ | $0.7179 \pm 0.0015$ | $0.7247 \pm 0.0006$ |
| Ours: GP pretraining only | $0.6881 \pm 0.0089$ | $0.7029 \pm 0.0038$ | $0.7152 \pm 0.0016$ | $0.7166 \pm 0.0008$ | $0.7185 \pm 0.0009$ | $0.7245 \pm 0.0008$ |
| PepCABO: Multi-RNC only | $0.6810 \pm 0.0116$ | $0.6974 \pm 0.0045$ | $0.7136 \pm 0.0022$ | $0.7176 \pm 0.0010$ | $0.7205 \pm 0.0011$ | $0.7259 \pm 0.0006$ |
| PepCABO: Full w/o $\mathcal{L}^{m}_{\mathrm{MRNC}}$ | $0.6893 \pm 0.0074$ | $0.7020 \pm 0.0040$ | $0.7159 \pm 0.0022$ | $0.7176 \pm 0.0009$ | $0.7197 \pm 0.0007$ | $0.7258 \pm 0.0006$ |
| PepCABO (Full model) | $0.6904 \pm 0.0079$ | $0.7037 \pm 0.0040$ | $0.7174 \pm 0.0016$ | $0.7188 \pm 0.0008$ | $0.7216 \pm 0.0009$ | $0.7262 \pm 0.0008$ |
| **Guided Initialization** | | | | | | |
| PepCABO: GP pretraining only | $0.6964 \pm 0.0047$ | $0.7025 \pm 0.0044$ | $0.7145 \pm 0.0027$ | $0.7173 \pm 0.0006$ | $0.7197 \pm 0.0009$ | $0.7256 \pm 0.0007$ |
| PepCABO: Multi-RNC only | $0.7166 \pm 0.0008$ | $0.7174 \pm 0.0009$ | $0.7218 \pm 0.0013$ | $0.7226 \pm 0.0004$ | $0.7235 \pm 0.0004$ | $0.7260 \pm 0.0007$ |
| PepCABO: Full w/o $\mathcal{L}^{m}_{\mathrm{MRNC}}$ | $0.7170 \pm 0.0008$ | $0.7180 \pm 0.0010$ | $0.7217 \pm 0.0010$ | $0.7223 \pm 0.0006$ | $0.7229 \pm 0.0006$ | $0.7264 \pm 0.0005$ |
| PepCABO: Full | $\mathbf{0.7173 \pm 0.0009}$ | $\mathbf{0.7179 \pm 0.0008}$ | $\mathbf{0.7221 \pm 0.0008}$ | $\mathbf{0.7231 \pm 0.0003}$ | $\mathbf{0.7239 \pm 0.0004}$ | $\mathbf{0.7266 \pm 0.0005}$ |

evaluated peptides per step. In this setting, the end-to-end update was performed at the beginning of each step (except the first step). Since the number of optimization steps and end-to-end updates is small, we ensured a fair comparison by requiring each optimization run to use the same number of steps. Therefore, we enforced the evaluation of precisely 20 new peptides at each step to maintain a fixed and controlled evaluation schedule. If an insufficient number of new candidates was found, we increased the radius of the trust-region-based search and reset it to its previous value at the start of the step after identifying enough new samples.

***PepPPO* Evaluation Protocol.** For *PepPPO*, optimization was conducted over 100 episodes with 8 mutations per episode in the low-budget setting. The maximum mutation length (8 steps) was chosen following Chen et al. [2023]. To determine the best peptide, the sequence at the end of each episode was evaluated. In the high-budget setting, candidates were assessed over 1000 episodes with the same maximum number of mutations per episode.

#### 5 Abliation Study

Table 2 contains an ablation study comparing the InvBO baseline to variants of our method. We evaluate (i) GP pretraining on the dual-VAE latent space without Multi-RNC, (ii) Multi-RNC alignment without GP pretraining, and (iii) the full model while omitting the contrastive loss term $\mathcal{L}^{m}_{\mathrm{MRNC}}$ during the optimization stage. Across both low- and high-budget settings, each component contributes to performance, and the full model achieves the strongest overall results, particularly under guided initialization. Under random initialization, both GP pretraining and Multi-RNC improve AUOC and final best-found affinity over InvBO, indicating complementary effects of surrogate calibration and latent alignment. Multi-RNC yields larger gains in the high-budget regime, suggesting that structured latent geometry becomes increasingly beneficial as exploration expands. Their combination consistently achieves the strongest performance, with improvements further amplified under guided initialization.

#### 6 Modified MRNC Loss for Mixed Data

Experimental binding affinity data consist of two types: quantitative and qualitative. In the former, an exact binding affinity value is assigned to each peptide–allele pair. In the latter, if the data originate from affinity measurements, only an upper or lower bound is provided instead of an exact value. If the data come from ligand evidence, no numerical affinity value is assigned. Following MHCflurry, we assign a lower bound of 0.574 (corresponding to 100 nM on the original scale) to such evidence.

We define $\mathcal{E}(\mathbf{x}_m), \mathcal{U}(\mathbf{x}_m)$, and $\mathcal{B}(\mathbf{x}_m)$ as the sets of peptides with exact measurements, upper-bound measurements, and lower-bound measurements, respectively, for a given MHC allele $\mathbf{x}_m$, with all values expressed in the log-transformed space. Hereafter, we treat the affinity value $f(\mathbf{x}_p, \mathbf{x}_m)$ for peptides in $\mathcal{U}(\mathbf{x}_m)$ and $\mathcal{B}(\mathbf{x}_m)$ as equal to their corresponding bound.

We now define the contrastive set for a given peptide–allele anchor pair $(\mathbf{x}_m, \mathbf{x}_p)$ only when $\mathbf{x}_p \in \mathcal{E}(\mathbf{x}_m) \cup \mathcal{B}(\mathbf{x}_m)$, as follows:

$$\mathcal{S}(\mathbf{x}_p, \mathbf{x}_m) := \{\mathbf{x}_{\hat{p}} \mid \mathbf{x}_{\hat{p}} \in \mathcal{E}(\mathbf{x}_m) \cup \mathcal{U}(\mathbf{x}_m),\ y_{\hat{p},m} \leq y_{p,m}\}.$$

Using the same negative Euclidean distance similarly, the Multi-RNC for experimental values becomes:

$$\mathcal{L}^*_{\text{MRNC}} = \frac{1}{N} \sum_{\mathbf{x}_m \in \mathcal{X}_m} \sum_{\mathbf{x}_p \in \mathcal{E}(\mathbf{x}_m) \cup \mathcal{B}(\mathbf{x}_m)} - \log \frac{\exp\big(\text{sim}(\mathbf{x}_p, \mathbf{x}_m)/\tau\big)}{\sum_{\mathbf{x}_{\hat{p}} \in \mathcal{S}(\mathbf{x}_p, \mathbf{x}_m)} \exp\big(\text{sim}(\mathbf{x}_{\hat{p}}, \mathbf{x}_m)/\tau\big)}. \tag{15}$$

### 7 Initialization Methods in Experimental Setting

Let $\mathcal{E}(\mathbf{x}_m)$ denote the set of peptides for which quantitative affinity measurements are available for a given MHC allele $\mathbf{x}_m$. To implement random initialization in the experimental setting, we randomly sample from all available peptides with quantitative binding affinity measurements for each of the 12 held-out alleles.

For guided initialization, we first pretrain the model as described in the manuscript section 2.2.3, with two modifications: (i) we use only quantitative experimental values to compute the surrogate loss, and (ii) for alignment, we apply $\mathcal{L}^*_{\text{MRNC}}$ as described in the previous section instead of the standard $\mathcal{L}_{\text{MRNC}}$. Applying the initialization method described in manuscript section 2.2.4 may result in peptides with $\mathbf{x}_p \notin \mathcal{E}(\mathbf{x}_m)$, which would require *in vitro* measurement. To mimic this method as closely as possible without generating unseen peptides, we instead rank peptides within $\mathcal{E}(\mathbf{x}_m)$ based on their Euclidean distance to the target MHC allele in the latent space and select the closest 10%. Finally, using Thompson sampling, we choose the best 20 candidates.

### References


Ziqi Chen, Baoyi Zhang, Hongyu Guo, Prashant Emani, Trevor Clancy, Chongming Jiang, Mark Gerstein, Xia Ning, Chao Cheng, and Martin Renqiang Min. Binding peptide generation for mhc class i proteins with deep reinforcement learning. *Bioinformatics*, 39(2):btad055, 01 2023. ISSN 1367-4811. doi: 10.1093/bioinformatics/btad055. URL `https://doi.org/10.1093/bioinformatics/btad055`.

Seunghun Lee, Jaewon Chu, Sihyeon Kim, Juyeon Ko, and Hyunwoo J. Kim. Advancing bayesian optimization via learning correlated latent space. In *Proceedings of the 37th International Conference on Neural Information Processing Systems*, NIPS '23, Red Hook, NY, USA, 2023. Curran Associates Inc.

Timothy J. O'Donnell, Alex Rubinsteyn, and Uri Laserson. Mhcflurry 2.0: Improved pan-allele prediction of mhc class i-presented peptides by incorporating antigen processing. *Cell Systems*, 11(1):42–48.e7, 2020. ISSN 2405-4712. doi: https://doi.org/10.1016/j.cels.2020.06.010. URL `https://www.sciencedirect.com/science/article/pii/S2405471220302398`.
